# Anxiolytic effects of chronic intranasal oxytocin on neural responses to threat are dose-frequency dependent

**DOI:** 10.1101/2021.04.20.440539

**Authors:** Juan Kou, Yingying Zhang, Feng Zhou, Zhao Gao, Shuxia Yao, Weihua Zhao, Hong Li, Yi Lei, Shan Gao, Keith M. Kendrick, Benjamin Becker

## Abstract

Anxiety disorders are prevalent psychiatric conditions characterized by exaggerated anxious arousal and threat reactivity. Animal and human studies suggest an anxiolytic potential of the neuropeptide oxytocin (OT), yet, while a clinical application will require chronic administration protocols previous studies in humans have exclusively focused on single-dose (acute) intranasal OT effects. We aimed at determining whether the anxiolytic effects of OT are maintained with repeated (chronic) administration or are influenced by dose frequency and trait anxiety. A double-blind randomized, placebo-controlled pharmaco-fMRI trial (n=147) determined acute (single-dose) as well as chronic effects of two different dose frequencies of OT (OT administered daily for 5 days or every other day) on emotional reactivity in healthy subjects with high versus low trait anxiety. OT produced valence, dose frequency and trait anxiety specific effects, such that the low-frequency (intermittand) chronic dosage specifically attenuated neural reactivity in amygdala-insula-prefrontal regions in high anxious subjects in response to threatening but not positive stimuli. The present trial provides evidence that low dose frequency chronic oxytocin nasal spray has the potential to alleviate exaggerated neural threat reactivity in subjects with elevated anxiety levels underscoring a treatment potential for anxiety disorders.

## Introduction

Anxiety is an evolutionary conserved adaptive response to uncertain future threats accompanied by strong negative emotional feelings (Sussman et al., 2016). Exaggerated anxious arousal and threat reactivity represent key symptoms of anxiety disorders, which - with a lifetime prevalence of 16% - are highly prevalent mental disorders (Kessler et al., 2009). However, current interventions for elevated anxiety are characterized by moderate response rates and often achieve only partial remission of symptoms (Blanco et al., 2013; Carpenter et al., 2018; Ipser et al., 2015). Translational approaches thus aim at the development of novel pharmacological strategies that specifically target underlying neurobiological mechanisms (Neumann & Slattery, 2016; Singewald et al., 2015; Zhou et al., 2019).

Anxiety engages a broad network of subcortical and cortical systems, with amygdala, insula and prefrontal (PFC) regions being among the most consistently identified (Bishop et al., 2004; Calder et al., 2011; Calhoon & Tye, 2015; Etkin & Wager, 2007; Larson et al., 2005; Robinson et al., 2019; Tovote et al., 2015; Xu et al., 2021; Zhao et al., 2019) in a network mediating arousal, alertness and vigilance (Sadaghiani & D’Esposito, 2015). Neuroimaging meta-analyses and treatment evaluation studies have demonstrated that patients with anxiety disorders exhibit exaggerated reactivity in this network in response to threat-signaling and negative emotional stimuli (Chavanne & Robinson, 2020; Etkin & Wager, 2007; Xu et al., 2021), while some functional dysregulations in the amygdala-insula-prefrontal circuitry normalize with successful intervention (Brehl et al., 2020; De Cagna et al., 2019; Goossens et al., 2007; S. M. Gorka et al., 2019; Ipser et al., 2015; Spengler et al., 2017).

Current translational perspectives propose an important regulatory role of the hypothalamic neuropeptide oxytocin (OT) in anxiety- and fear-related domains. Endogenous OT release critically regulates the fear response of the amygdala (Knobloch et al., 2012) and the formation of fear memory (Hasan et al., 2019), while intracerebroventricular administration of OT reduces anxiety (Ayers et al., 2011) and facilitates fear extinction (Triana-Del Río et al., 2019; Toth et al., 2012) in rodents. Potential anxiolytic properties of OT have been documented by proof-of-concept studies in healthy individuals reporting that intranasal administration of a single dose (Lee et al., 2020; Quintana et al., 2020) attenuates amygdala responses during exposure and anticipation of threatening stimuli (Kirsch, 2005; Xin et al., 2020), facilitates fear extinction learning via enhancing prefrontal activity (Eckstein et al., 2015) and modulates insula activity during approach and attention towards salient emotional stimuli (Striepens et al., 2012; Yao, Becker, et al., 2018).

Based on these findings and overarching reviews emphasizing the anxiolytic properties of OT (Gottschalk & Domschke, 2018; Neumann & Slattery, 2016), initial clinical trials explored its therapeutic potential in patients with anxiety disorders. These studies generally documented a promising potential of OT to attenuate core neuro-functional dysregulations in anxiety disorders, including an attenuation of amygdala and PFC hyper-reactivity as well as enhanced functional connectivity between the amygdala with prefrontal and insular regions during exposure to negative stimuli in patients with generalized (social) anxiety disorder (Gorka et al., 2015; Labuschagne et al., 2010, 2012; De Cagna et al., 2019). However, studies to date examined potential anxiolytic effects following a single dose of intranasal OT, while therapeutic application will require long-term (chronic) administration protocols and maintenance of anxiolytic effects (De Cagna et al., 2019).

While there is consistent evidence for an anxiolytic potential of a single dose of OT, animal studies examining long-term treatment protocols have reported inconsistent or even anxiogenic effects. Chronic administration of high doses produced anxiogenic effects in rodents (Peters et al., 2014;Winter et al., 2020) which may reflect a consequence of chronic exposure-induced regional-specific downregulation or desensitization of endogenous OT receptors (Pisansky et al., 2017). On the other hand, the chronic administration of lower doses attenuated anxiety-related behavior in high trait but not low trait anxiety animals (Slattery & Neumann, 2010). These findings suggest differential, dose- and trait anxiety-dependent effects of acute versus chronic OT administration on anxiety. Furthermore, the translational importance of these findings is documented by a recent study reporting increased anxiety-like behavior in male juvenile non-human primates after chronic OT administration (Razo et al., 2020) and another study in men demonstrating that a single dose of intranasal OT reduced amygdala reactivity towards facial stimuli, while neural and behavioral responses to threatening facial expressions were abolished after five days of daily chronic treatment but maintained when it was administered every other day (Kou et al., 2020).

Taken together, these findings indicate differential or even opposing acute versus chronic effects of intranasal OT on anxiety-related domains which are of critical relevance for the clinical application of OT in anxiety disorders. Against this background the present randomized placebo-controlled pharmaco-fMRI trial examined acute and dose frequency-dependent chronic effects of OT during implicit processing of negative and positive emotional scenes. Given that previous studies suggest acute and chronic effects of OT are influenced by pre-treatment differences in trait anxiety (Alvares et al., 2012; Ayers et al., 2011; Luo et al., 2017; Schumacher et al., 2018; Slattery & Neumann, 2010) and increase the translation relevance of findings to populations with elevated anxiety, individuals pre-screened for high (HTA) and low (LTA) levels of trait anxiety were enrolled. Based on the previous literature we hypothesized that: (1) a single dose of OT would specifically attenuate neural and behavioral reactivity towards negative emotional stimuli, particularly in HTA subjects, while (2) daily chronic OT treatment for five days, but not low dosage treatment given every other day, would abolish its anxiolytic effects.

## Methods and Materials

### Participants

The main aim of the present randomized double-blind placebo controlled pharmacological fMRI trial was to determine whether acute and chronic effects of OT on emotional processing differ in subjects with high versus low trait anxiety. From a total of n = 917 subjects that underwent screening 147 male participants (age range 18-27years) with high or low trait anxiety were enrolled in the present study. Based on previous studies (Ironside et al., 2019) and local trait anxiety distribution (details see SI) total trait anxiety scores as assessed by the Spielberger State-Trait Anxiety Inventory (STAI) (Kvaal et al., 2005) of ≥45 were considered as the cut-off for the high anxiety group, whereas scores ≤ 35 were considered as the cut-off for the low anxiety group (for trait anxiety levels see **Table 1**, details **supplementary information**).

**Table1.** Anxiety and depression scores in all subgroups

| Questionnaires | High anxiety (Mean±SD) |  |  |  | Low anxiety (Mean±SD) |  |  |  |
| --- | --- | --- | --- | --- | --- | --- | --- | --- |
|  | PLC(n=24) | OT<sub>3</sub>(n=22) | OT<sub>5</sub>(n=24) | *p* | PLC(n=26) | OT<sub>3</sub>(n=25) | OT<sub>5</sub>(n=23) | *p* |
| TAI | 51.5±5.0 | 50.5±5.9 | 48.7±3.4 | 0.14 | 30.1±4.5 | 30.9±3.5 | 30.4±4.4 | 0.81 |
| SAI | 45.3±9.9 | 43.8±8.7 | 44.6±6.0 | 0.83 | 29.5±6.0 | 28.0±5.5 | 31.1±5.7 | 0.18 |
| BDI | 14.5±8.2 | 14.9±7.7 | 12.1±8.4 | 0.46 | 3.5±4.2 | 3.3±3.6 | 4.4±5.0 | 0.62 |

Additional exclusion criteria were: (1) history/current psychiatric or neurological disorder, (2) current or regular use of psychotropic substances, and (3) MRI or OT contraindications. To account for sexual dimorphic effects of OT (e.g. Gao et al., 2016; Lieberz et al., 2019; Ma et al., 2018) and menstrual cycle dependent fluctuations in endogenous OT levels (Engel, Klusmann, et al., 2019; Engel, Laufer, et al., 2019) the study focused on male participants (for similar approach see also Xin et al., 2020.; Yao, Becker, et al., 2018). Data from n = 3 subjects were excluded due to excessive head motion (3mm translation or 3° rotation or mean frame-wise displacement > 0.5mm).

Written informed consent was obtained from all participants and subjects received monetary compensation. The study was approved by the local ethics committee (University of Electronic Science and Technology of China) and pre-registered on Clinical Trials.gov (https://clinicaltrials.gov/ct2/show/NCT03085654).

### Design and experimental protocols

Eligible subjects were enrolled in a double-blind, randomized, placebo-controlled, between-subject pharmacological trial with fMRI assessments following acute (first day of treatment) and chronic (fifth day of treatment) OT administration. Subjects in the high or low trait anxiety groups were randomly assigned to three different intranasal treatment protocols (PLC, OT_3_ and OT_5_ groups) (see **Figure1**, Consort flow diagram - single daily dose on five consecutive days of placebo (PLC) or oxytocin (OT_5_), or interleaved OT and PLC (OT on days 1, 3, 5, PLC on days 2, 4; OT_3_) To promote compliance with the standardized intranasal treatment protocols (Guastella et al., 2013) subjects were required to come to the laboratory during five consecutive days to self-administer the nasal spray under supervision. Following treatment administration on day 1 and day 5, all subjects underwent an emotion processing fMRI paradigm to determine trait anxiety-dependent influences on the acute (day 1) and chronic (day 1 compared to day 5) effects of OT. Details of the experimental procedures are provided in **Figure 2**.

**Figure 1.**
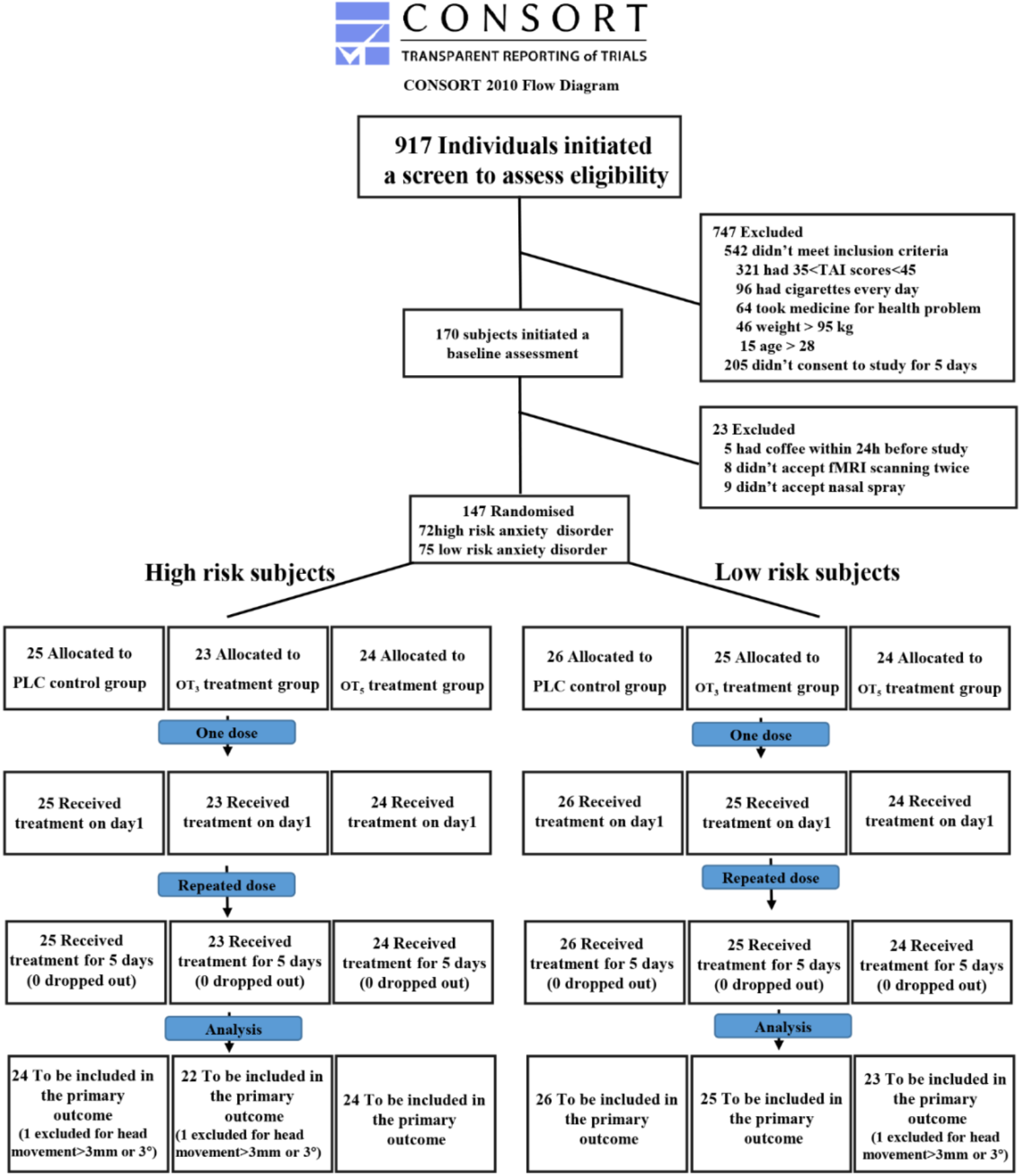
CONSORT flow diagram of the clinical trial

**Figure 2.**
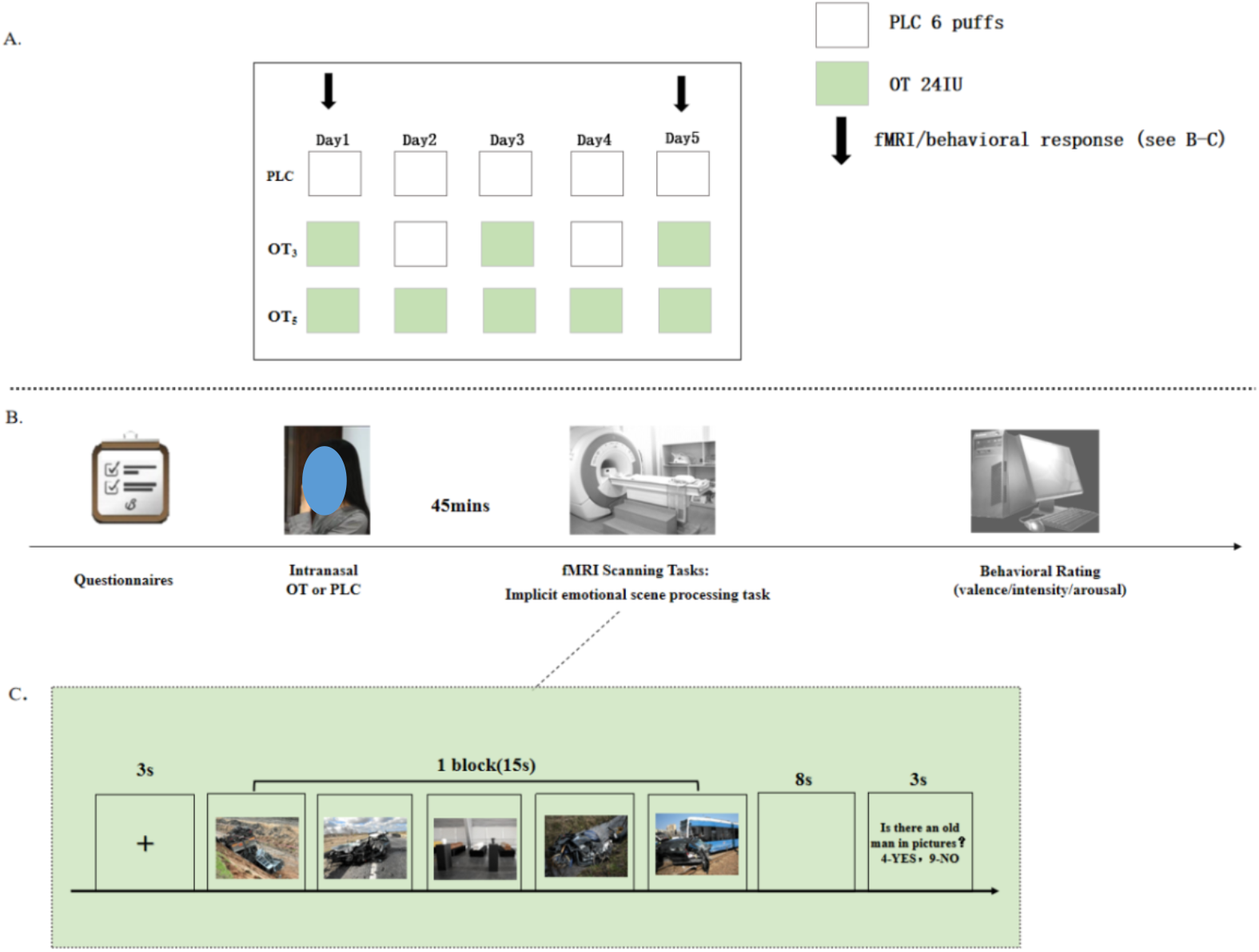
Treatment protocol. (A) Participants were randomly assigned to intranasal oxytocin (OT) for 5 days with 24 IU per day (OT_5_ group) or to receive OT or placebo nasal spray on alternate days during the 5 days (OT on the 1^st^, 3^rd^ and 5^th^ day), 24 IU per day (OT_3_ group). The placebo group received placebo nasal spray for 5 consecutive days (PLC group); (B) Experimental timeline and (C) emotion processing fMRI paradigm, (D) post fMRI behavioral ratings. Note: For the preprint version the face in Figure 2b was overlayed with a blue shape.

A dose of 24 International Units (IU) of OT was administered based on previous studies reporting its acute (Spengler et al., 2017) and dose-frequency dependent(Kou et al., 2020) neural effects. In line with recent pharmacodynamic studies reporting brain activity changes around 45 minutes after administration (Paloyelis et al., 2016) experiments started 45 minutes after intranasal administration. OT and PLC sprays were both supplied by Sichuan Meike Pharmaceutical Co. Ltd, Sichuan, China in identical dispenser bottles containing identical ingredients (glycerine and sodium chloride) other than OT.

Based on the high co-morbidity between anxiety and depression, and previous studies reporting that subclinical levels of depression influence effects of OT (Eckstein et al., 2017) levels of depression were assessed before the first treatment using the Beck Depression Inventory (BDI II) (Beck et al., 1988). Although there were no significant differences between the OT and PLC groups in either anxiety group (see **Table 1**), high trait anxiety subjects generally reported higher levels of depressive symptoms compared to the low anxiety group. Depression was therefore included as a covariate in all subsequent analyses.

### fMRI emotional processing paradigm

During the emotion processing fMRI paradigm a total of 180 emotional scene stimuli from the Nencki Affective Picture System (Marchewka et al., 2014) with negative, positive and neutral emotional content (n = 30 per category) were displayed. To control for stimulus-dependent repetition effects two sets of stimuli with 90 stimuli each were used and the order of the stimuli sets was randomized across subjects. Stimuli sets were matched for valence and arousal (details see **Table S1**).

An implicit emotional processing paradigm with a blocked design encompassed a total of 18 blocks (6 blocks per emotional category) containing 5 condition-specific stimuli per block. Each stimulus was presented for 3s and blocks interspersed by a low-level baseline with a fixation jitter presented for 12-16 seconds. To ensure attentive processing each block was followed by a question referring to the content of the pictures shown (e.g. ‘was an old man shown?’) with subjects responding by button press (see **Figure 2C**). To assess behavioral subjects rated arousal, intensity and valence of stimuli on 9-point Likert scales after each MRI acquisition (see **Figure 2**).

### Image acquisition

Blood oxygenation level-dependent contrast functional time series were acquired using standard sequences on a 3T GE MR750 system. Functional images for task-based fMRI were acquired using an EPI sequence (TR=2000ms, echo time=30ms, flip angle=90°, FOV=240×240mm, voxel size=3.75×3.75×4 mm, resolution=64×64, number of slices=39). T1-weighted anatomical images were additionally acquired to improve normalization of the functional images (TR=6ms, echo time=2ms, flip angle=9°, FOV=256×256mm, voxel size=1×1×1mm, number of slices=156).

### Primary outcomes and analysis plan

#### Neural level - emotional brain processing

Functional MRI data was preprocessed and analyzed using SPM12 (Friston et al., 1994) (Statistical Parametric Mapping; http://www.fil.ion.ucl.ac.uk/spm). The first 5 volumes were discarded to allow T1 equilibration and the remaining images were corrected for temporal slice acquisition differences, realigned to the first volume, and unwarped to correct for nonlinear distortions related to head motion (six-parameter rigid body algorithm). Tissue segmentation and bias-correction were applied to the high-resolution structural images. After co-registration of the functional time-series with the skull-stripped anatomical images the transformation matrix was applied to normalize the functional images to MNI standard space with a voxel size resolution of 3×3×3mm. Normalized images were spatially smoothed using a Gaussian kernel with full-width at half-maximum (FWHM) of 5mm. Participants with head motion exceeding 3mm translation or 3° rotation and mean frame-wise displacement (FD) > 0.5mm were excluded.

A random effects general linear model (GLM) analysis was employed for statistical analyses. First level GLMs for the fMRI data included separate regressors for the three emotional conditions (negative, positive and neutral emotional stimuli) and the period during which subjects had to indicate their response to the question following each block. A high-pass filter of 128 seconds was applied to remove low frequency drifts. Head motion parameters were included (six-parameter rigid body algorithm) as covariates. Contrasts of interest to examine the acute ([negative-neutral] _day1_, [positive-neutral] _day1_) and chronic effects of OT ([negative-neutral] _day5_, [positive-neutral] _day5_) were produced on the individual level and subjected to second level analysis.

In line with the major aims of the study a second level model was designed to determine interactive effects between trait anxiety and repeated dosage regimes by means of mixed ANOVA models with the factors treatment groups (OT_3_, OT_5_, PLC) × emotional valence (negative minus neutral, positive minus neutral) × time point (day1, day5) × trait anxiety (high anxiety, low anxiety) encompassing data from both days. Multiple comparisons were corrected on the whole-brain level by employing Family-wise error (FWE) correction with *p* < 0.01 at the cluster level (cluster forming threshold p < .001, minimum cluster size > 180 voxels). To disentangle significant interaction effects, post-hoc analysis were performed on extracted parameter estimates using independent atlas-based masks for the identified brain regions (Brainnetome atlas, Fan et al., 2016). In line with the primary aim of the study post-hoc pair-wise tests focused on comparing the experimental groups on each day and comparison of changes over the course of chronic administration between the experimental groups. To further control for potential effects of differences in depression post hoc pair-wise comparisons included BDI scores as a covariate. Bonferroni correction was conducted for post-hoc pair-wise comparisons.

#### Behavioral level – subjective emotional experience

Effects on the behavioral level (valence, intensity and arousal ratings) were examined by four way ANOVA models for acute and chronic effects with the factors treatment groups (OT_3_, OT_5_, PLC) × emotional valence (negative minus neutral, positive minus neutral) × time point (day1, day5) × trait anxiety (high anxiety, low anxiety) encompassing data from both days. Post-hoc comparisons with Bonferroni correction were conducted for significant interactions.

## Results

### Acute and chronic effects of OT and modulation by dose frequency and anxiety

Whole-brain voxel-wise mixed ANOVAs including the factors treatment, trait anxiety, emotional valence and time point, revealed a significant interaction effect in the left amygdala (peak MNIxyz = [−18,−4,−22], *F*_(2,138)_ = 9.49, *P*clusterFWE < 0.007, k = 197), right anterior insula (peak MNIxyz = [33, 5, −10], *F*_(2,138)_ = 11.91, *P*clusterFWE < 0.001, k = 340), and left dorsolateral prefrontal cortex (dlPFC) (peak MNIxyz = [−30, 11, 41], *F*_(2,138)_ = 15.71, *P*clusterFWE < 0.001, k = 521), as well as brain stem, inferior parietal lobule and middle cingulate cortex regions (see **Figure3**). Post-hoc analysis of the extracted estimates from the left amygdala, right anterior insula (masks from the Brainnetome (Fan et al., 2016)) and left dlPFC (Brodmann areas 9 and 46 (Mylius et al., 2013)) revealed that a single dose of OT did not affect activity in these regions (*ps* _bonferroni_ > 0.337), while examination of longitudinal changes (day 5 minus day 1) revealed differences between the PLC and OT_3_ high trait anxiety groups in response to negative stimuli (*p* _bonferroni_ amygdala = 0.033, dlPFC = 0.02; insula = 0.049), reflecting increasing activity in the PLC but decreasing activity in the OT_3_ high trait anxiety groups. No changes were observed in the OT_5_ group (*ps* > 0.06). In line with the longitudinal changes, post-hoc analysis on the day 5 data revealed that activation of the left amygdala (*p* _bonferroni_ =0.013), left dlPFC (*p* _bonferroni_ =0.007) and right anterior insula (*p* =0.031) in response to negative stimuli was significantly lower in the OT_3_ high anxiety trait group as compared to the PLC one. No other post hoc comparisons of treatment on day 5 reached significance (*ps* > 0.06) (see **Figure 3**). No significant treatment effects for the positive stimuli were observed (*ps* _bonferroni_ > 0.09), suggesting a negative processing-specific effect. No significant main effects of treatment or trait anxiety were observed, arguing against unspecific effects of these factors.

**Figure 3.**
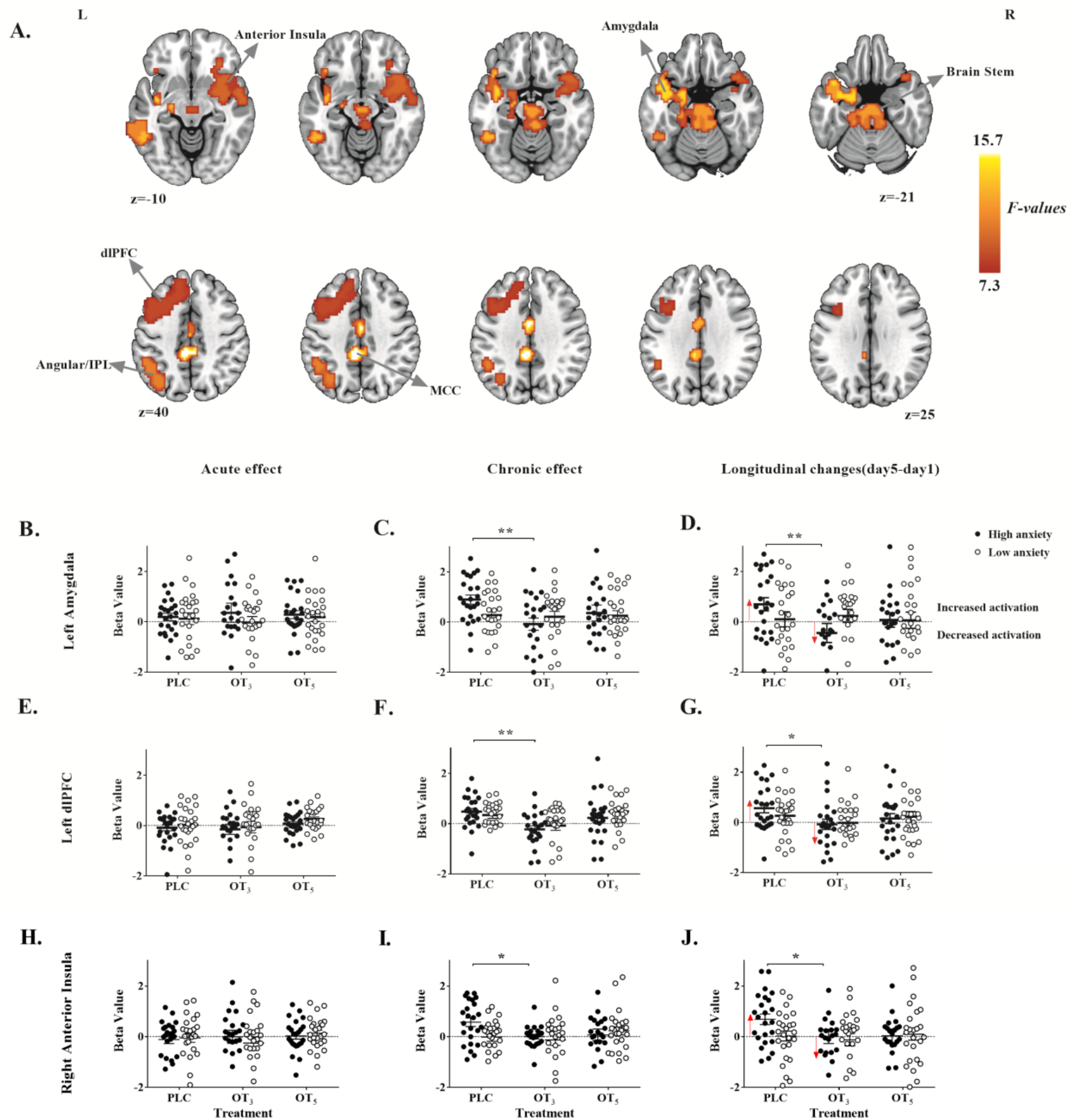
Longitudinal low dose frequency intranasal OT decreased activation of left amygdala and left dlPFC to negative scenes. (A) Axial slices showing the location of the 4 way-interaction effect (treatment groups× emotional valence× time point (day1, day5) × trait anxiety) displayed as F map (*p*< 0.001, Cluster size≥180) with significant clusters located in the left amygdala, right insula, brain stem, left dlPFC (dorsolateral prefrontal cortex), left IPL (inferior parietal lobule) and MCC (middle cingulate cortex). (B)-(D) Results of post-hoc analyses of the left amygdala (beta value extracted from an independent mask as defined by the Brainnetome atlas), and (E)-(G) post-hoc analyses of left dlPFC (beta value extracted by using mask from Brodmann areas 9 and 46 (Mylius et al., 2013)) to negative scenes. (H)-(J) Results of post-hoc analyses of the right anterior insula (beta values extracted from an independent mask as defined by the Brainnetome atlas) * p < 0.05, ** p < 0.01, two-tailed t-test. Bars indicate standard errors.

### Acute and chronic OT effects on the behavioral level

Mixed four-way ANOVAs with treatment (PLC, OT_3_ and OT_5_), trait anxiety (high trait anxiety, low trait anxiety), time point (day1, day5) and stimuli valence (negative-neutral, positive-neutral) as factors and emotional valence, intensity and arousal ratings as dependent variable were conducted. For emotional valence and arousal ratings, there was no significant four-way interaction effect (*ps*>0.084). Although the 4 way interaction effect was significant for intensity ratings (F_2, 137_=3.89, *p* = 0.023, η^2^p =0.054), pair-wise post hoc comparisons failed to reach statistical significance (*ps* > 0.113). Moreover, no main effects of treatment on emotional valence, intensity and arousal ratings reached statistical significance (ps>0.373).

## Discussion

Overall, our findings revealed that the chronic effects of OT on emotional brain activity vary as a function of both trait anxiety and dose frequency. Although a single dose of OT did not affect neural activity during processing of positive and negative emotional stimuli, administration of OT every other day (OT_3_) specifically decreased neural reactivity specifically towards negative stimuli in the amygdala, insula and dlPFC in high trait anxiety subjects, while the PLC-treated high trait anxiety group exhibited increased activity in these regions. In contrast, chronic daily administration of OT (OT_5_) did not affect neural activity in either of the trait anxiety groups. No effects on the behavioral level in terms of explicit subjective valence, intensity and arousal perception were observed.

### No evidence for acute anxiolytic effects of OT on neural reactivity

In contrast to our hypothesis and previous animal and human studies reporting effects of a single dose of OT on anxiety-related behavior and brain activation in response to negative social stimuli (Neumann & Slattery, 2016), the present study did not observe behavioral or neural effects in response to either positive or negative emotional scenes following single dose administration on day 1. Previous task-fMRI studies reporting attenuated neural reactivity in response to negative emotional stimuli in healthy male subjects following a single dose of OT - frequently interpreted as potential anxiolytic effects of OT ‒predominately employed explicitly social, primarily facial, stimuli (Kou et al., 2020; Kreuder et al., 2020; Quintana et al., 2016). In contrast, effects of OT on non-social stimuli or complex scenes have been less consistent, such that a single dose of OT increased anxiety to unpredictable threat (Grillon et al., 2013), did not affect neural reactivity in response to negative scenes (Xin et al., 2020) or increased insular reactivity in response to successfully encoded negative emotional scenes (Striepens et al., 2012). Moreover, effects of OT on neural threat processing have been linked to highly specific features such as threat signals from the eye region of faces (Kanat et al., 2015) or to specific threat processing instructions including approach versus avoidance(Radke et al., 2017; Yao, Zhao, et al., 2018). In line with the increasing evidence for highly context- and social-specific effects of OT (Bartz et al., 2011; Shamay-Tsoory & Abu-Akel, 2016; Xu et al., 2019) our findings do not support general and rather unspecific neuro-anxiolytic effects of OT during processing of complex social and non-social emotional stimuli.

### Chronic effects of OT depend on trait anxiety and dose frequency

In contrast to a lack of effects of a single dose of OT, valence, trait anxiety and dose frequency-dependent chronic effects were observed in insula-amygdala-prefrontal regions. Whereas repeated daily doses of OT did not affect neural processing in subjects with low trait anxiety, or in response to positive stimuli, intermittant administration every other day attenuated neural activation in the amygdala, the anterior insula and dlPFC in high trait anxiety subjects during processing of negative stimuli. Further examination of the high trait anxiety treatment groups revealed that whereas the OT_3_ group exhibited decreased neural reactivity, the PLC treated group exhibited increased activity while the OT_5_ group did not change at all over the treatment period. Recent meta-analyses demonstrated exaggerated activity in the amygdala and insula in patients with anxiety disorders and individuals with high trait anxiety, particularly in response to negative emotional material (Chavanne & Robinson, 2020; Etkin & Wager, 2007; Stein et al., 2007), with observed alterations being suggested to contribute to exaggerated threat reactivity and anxious arousal. With respect to neurofunctional alterations in the dlPFC both increased as well as decreased activity has been reported in anxiety disorder. Attenuated dlPFC activity during cognitive emotion regulation is interpreted to underlie deficient regulatory control, while increased activity during emotion perception may reflect enhanced attentional and salience processing of potentially threatening stimuli (Brehl et al., 2020; Bruce et al., 2013; Morgan et al., 1993; Quirk et al., 2003). Moreover, the anterior insula and lateral PFC have been suggested to play a role in several general domains associated with anxious arousal, including tonic alertness and arousal as well as threat specific domains such as expectation of predictable threat and threat vigilance (Ironside et al., 2016; Wheelock et al., 2014). Within this context, decreasing activity in these regions following a chronic intermittant dose of OT may reflect a potential anxiolytic rather than an anxiogenic potential of low-dose chronic OT in subjects with high trait anxiety showing decreasing alertness, threat reactivity and salience processing. In contrast to the OT-treated groups the PLC-treated high-, but not the low-trait anxiety group demonstrated increasing activity in these regions over the course of the chronic administration protocol. Given that the individuals in this group did not receive active treatment and increasing activity was specifically observed in the high trait anxiety group, interaction effects of trait anxiety with other factors of the design such as the repeated exposure to emotional stimuli may account for this effect. One possible explanation might be an elevated anticipatory anxiety in the high trait anxiety subjects for the repeated task, such that previous studies have demonstrated the anticipation of negative outcomes (eg., aversive pictures) recruits a neural network encompassing, among other regions, the amygdala, insula and dorsolateral prefrontal cortex (dlPFC)(Sarinopoulos et al., 2010; Straube et al., 2007) which may have promoted enhanced attentional salience of the negative stimuli.

A comparable pattern of effects involving trait anxiety and treatment was observed in other brain regions such as brain stem which is critically involved in the regulation of autonomic nervous system responses during exposure to stress and fear-inducing stimuli. In response to these stimuli both OT and vasopressin regulate sympathetic and parasympathetic pathways and mediate the autonomic stress response via oxytocinergic receptors in the brainstem(Freeman et al., 2016; Quintana & Guastella, 2020).

In contrast to the widespread neural effects of OT treatment in high anxiety individuals no effects of either treatment regimen were observed on the behavioral level. These divergent effects on the behavioral and neural levels may be explained in terms of differences in the brain systems and behavioral processes underlying implicit emotional versus explicit emotional processing, which vary in terms of the strengths of their association (Bargh, 1994; Nosek, 2007). Given that our major focus was to examine bottom-up and everyday emotional reactivity and that previous studies on the effects of OT on explicit, rather cognitive, emotional evaluations have been inconclusive (Tully et al., 2018) (Olivera-Pasilio & Dabrowska, 2020), we decided to employ an implicit processing paradigm. However, this may have come at the cost of a reduced sensitivity on the behavioral level.

Other limitations of the present study include the focus on male subjects, and although this is in line with other recent studies investigating kinetics and dosage effects of OT (Martins et al., 2020; Paloyelis et al., 2016; Spengler et al., 2017) the generalizability of the present findings to women needs to be established. To control for a range of unspecific factors the present study moreover employed a proof-of-concept design in high trait anxiety subjects and clinical utility of OT in patient populations with exaggerated anxiety remains to be evaluated.

Summarizing, the present results suggest valence, dose-frequency and trait anxiety dependent effects of chronic OT administration on a neural circuitry encompassing the amygdala, anterior insula and lateral prefrontal cortex. An intermittant dosage of chronic OT specifically decreased neural reactivity within this circuit in response to negative stimuli in subjects with high trait anxiety. An intermittant chronic dose regime may therefore have the clinical potential to alleviate exaggerated neural threat vigilance and reactivity.

## Acknowledgments

This study was supported by the National Key Research and Development Program of China (2018YFA0701400). Shenzhen-Hong Kong Institute of Brain Science-Shenzhen Fundamental Research Institutions (2019SHIBS0003).

