## Supplemental Information for "Anxiolytic effects of chronic intranasal oxytocin on neural responses to threat are dose-frequency dependent"

Supplementary Methods

Supplementary Table

Supplementary Figure

**Supplementary Methods**

Recruitment and selection of participants

In line with a previous study (Ironside et al., 2019) subjects with a trait anxiety scores ≥45 on the Spielberger State-Trait Anxiety Inventory (STAI) (Kvaal et al., 2005) were assigned to the high trait anxiety group. The cut-off score for the low-trait anxiety group (<35 STAI trait anxiety) was determined according to the distribution of trait anxiety in the local population. To this end a total of 917 subjects were screened with the STAI. 33% exhibited a score above 45 confirming the literature-based cut-off score for the high trait anxiety group. 33% of our subjects scored lower than 35 which as therefore determined as lower anxiety subgroup cut-off. The STAI scores in the local population were normal distributed (mean ± SD = 41.06±10.13) (see also Figure S1).

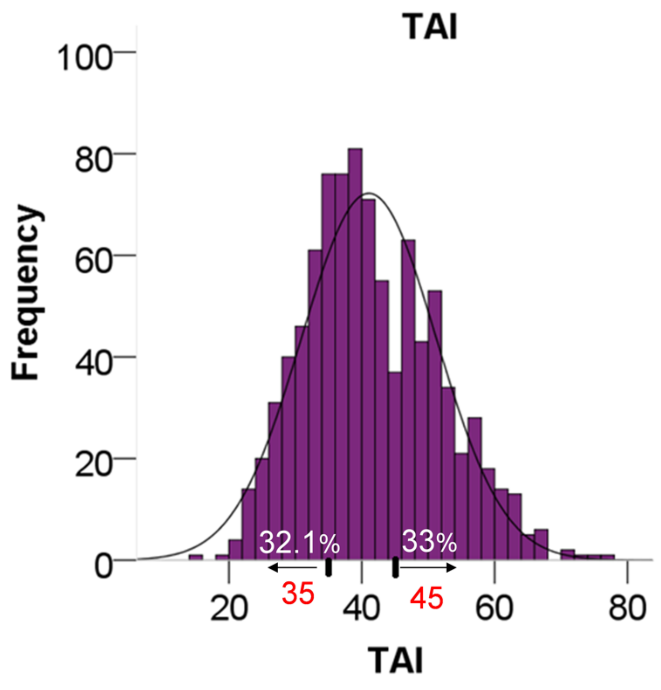


Figure S1 Distribution of trait anxiety scores (TAI) in the local population (n=917).

Table S1 Ratings for each of the two sets of different emotion stimuli used on the 1st day and 5th day in a counterbalanced design. There were no significant differences between the two sets for valence and arousal ratings.

| Valence | | | | | Arousal | | | |
|---|---|---|---|---|---|---|---|---|
| Emotional Categories | Set A On the 1st day | Set B On the 5th day | t | p | Set A On the 1st day | Set B On the 5th day | t | p |
| Positive | 7.41±0.46 | 7.40±0.49 | 0.09 | 0.93 | 4.00±0.88 | 4.02±0.98 | 0.17 | 0.87 |
| Neutral | 5.19±0.36 | 5.24±0.30 | 0.58 | 0.57 | 4.78±0.29 | 4.82±0.46 | 0.45 | 0.66 |
| Negative | 3.12±0.82 | 3.34±0.89 | 1.02 | 0.32 | 6.51±0.56 | 5.57±058 | 0.43 | 0.67 |
